# Discovery of BAZ2A Bromodomain Ligands

**DOI:** 10.1101/122317

**Authors:** Dimitrios Spiliotopoulos, Eike-Christian Wamhoff, Graziano Lolli, Christoph Rademacher, Amedeo Caflisch

## Abstract

The bromodomain adjacent to zinc finger domain protein 2A (BAZ2A) is implicated in aggressive prostate cancer. The BAZ2A bromodomain is a challenging target because of the shallow pocket of its natural ligand, the acetylated side chain of lysine. Here, we report the successful screening of a library of nearly 1500 small molecules by high-throughput docking and force field-based binding-energy evaluation. For seven of the 20 molecules selected *in silico*, evidence of binding to the BAZ2A bromodomain is provided by ligand-observed NMR spectroscopy. Two of these compounds show a favorable ligand efficiency of 0.42 kcal/mol per non-hydrogen atom in a competition-binding assay. The crystal structures of the BAZ2A bromodomain in complex with four fragment hits validate the predicted binding modes. The binding modes of compounds **1** and **3** are compatible with ligand growing for optimization of affinity for BAZ2A and selectivity against the close homologue BAZ2B.

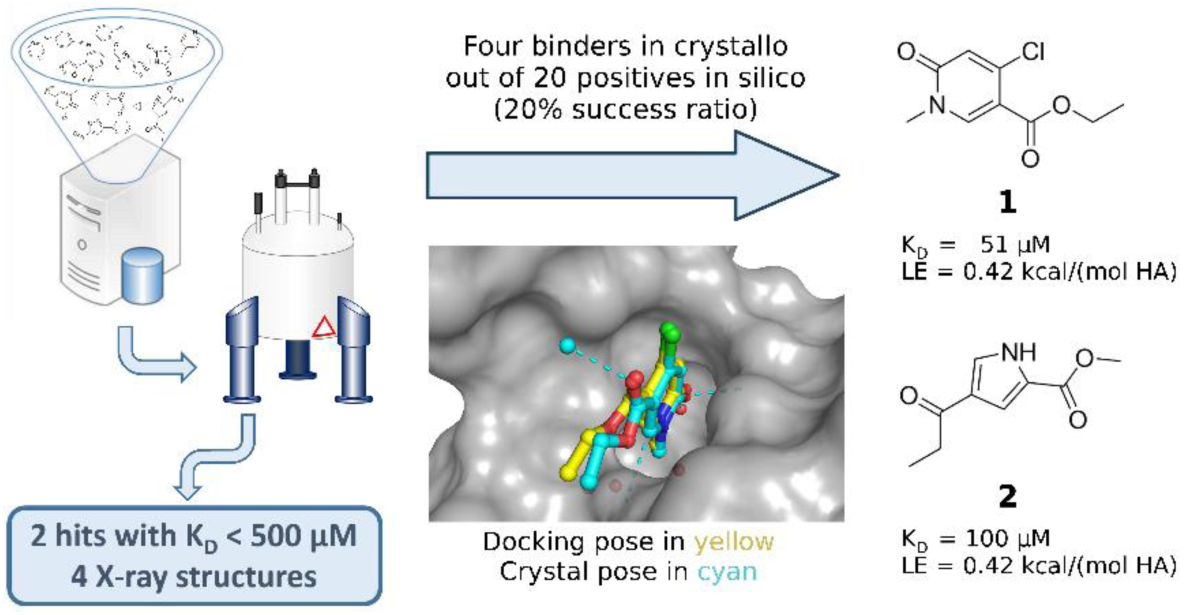

## INTRODUCTION

Bromodomains are epigenetic readers with a four helix-bundle topology.^1,2^ They recognize the acetylated side chain of lysine (Kac)^1^ as well as non-acetyl acyllysine modification, *e. g.*, propionyllysines, butyryllysines, and crotonyllysines^3^ in histones and other proteins.

The bromodomain-containing protein BAZ2A (bromodomain adjacent to zinc finger domain protein 2A) is involved in maintaining cell growth in prostate cancer and its overexpression is related to disease recurrence.^4^ The bromodomains of BAZ2A and BAZ2B (a bromodomain-containing protein similar to BAZ2A) are considered challenging to target by small molecules because of their shallow binding site.^5^ Only two potent inhibitors of the BAZ2A bromodomain have been reported as of today. Drouin et al. developed a potent BAZ2A/BAZ2B inhibitor starting from a double-digit micromolar hit. Two cycles of design and synthesis led to a nanomolar, selective BAZ2A/B inhibitor.^6^ In another study, an indolizine derivative reported in the literature as bromodomain and extra-terminal (BET) bromodomain inhibitor^7^ was identified by *in vitro* screening as single-digit micromolar binder of the BAZ2A bromodomain. Structure-based design efforts led to the generation of a submicromolar ligand of the BAZ2A/B bromodomains capable of engaging a green-fluorescent protein–BAZ2A fusion construct in the FRAP assay.^8^

Here, we report on the identification of BAZ2A bromodomain ligands by *in silico* screening of a library of nearly 1500 small molecules followed by biophysical validation of the top 20 compounds (**1**–**4**, Table 1; **5**–**20**, Table S1). Seven of the 20 compounds predicted by docking and scoring were confirmed as binders by ligand-based NMR spectroscopy. Two of them (compounds **1** and **2**, Table 1) show micromolar affinity and favorable ligand efficiency for the bromodomains of BAZ2A and BAZ2B in an *in vitro* assay. The crystal structures of the complexes of compounds **1**–**4** in the BAZ2A bromodomain validate the predicted binding modes. Moreover, these crystal structures allow us to analyze the binding modes in two bromodomains with very similar Kac binding site, and to exploit minor differences for the design of selective BAZ2A inhibitors.

**Table 1.**
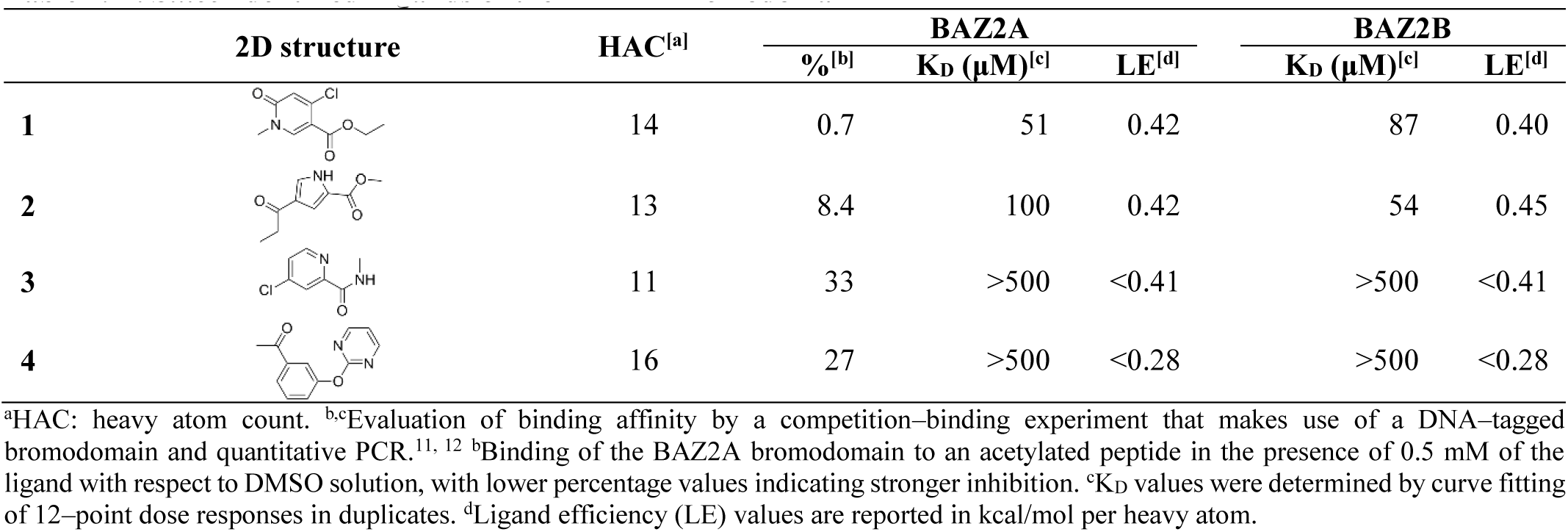
*In Silico* Identified Ligands of the BAZ2A Bromodomain

|  | 2D structure | HAC<sup>[a]</sup> | BAZ2A |  |  | BAZ2B |  |
| --- | --- | --- | --- | --- | --- | --- | --- |
|  |  |  | %<sup>[b]</sup> | K<sub>D</sub> (μM)<sup>[c]</sup> | LE<sup>[d]</sup> | K<sub>D</sub> (μM)<sup>[c]</sup> | LE<sup>[d]</sup> |
| 1 |  | 14 | 0.7 | 51 | 0.42 | 87 | 0.40 |
| 2 |  | 13 | 8.4 | 100 | 0.42 | 54 | 0.45 |
| 3 |  | 11 | 33 | >500 | <0.41 | >500 | <0.41 |
| 4 |  | 16 | 27 | >500 | <0.28 | >500 | <0.28 |

<sup>a</sup>HAC: heavy atom count. <sup>b</sup>cEvaluation of binding affinity by a competition-binding experiment that makes use of a DNA-tagged bromodomain and quantitative PCR.<sup>11, 12</sup> <sup>b</sup>Binding of the BAZ2A bromodomain to an acetylated peptide in the presence of 0.5 mM of the ligand with respect to DMSO solution, with lower percentage values indicating stronger inhibition. <sup>c</sup>K<sub>D</sub> values were determined by curve fitting of 12-point dose responses in duplicates. <sup>d</sup>Ligand efficiency (LE) values are reported in kcal/mol per heavy atom.

## RESULTS AND DISCUSSION

### *In silico* Screening

A library of 1413 small molecules (Figure S1) was docked into the Kac binding site of the BAZ2A bromodomain using the program SEED (details in Methods section). These molecules represent an existing and thoroughly maintained library which is readily available for screening in the laboratory of one of the authors (C.R.). The resulting poses were ranked by consensus scoring using the van der Waals and electrostatic energy contributions taking into account electrostatic desolvation effects in the continuum dielectric approximation (Figure S2).^9,10^ After the energy-based ranking (cf. Experimental Section), 20 small molecules were selected for validation by ligand-based NMR spectroscopy (Figure 1, Table S1).

**Figure 1.**
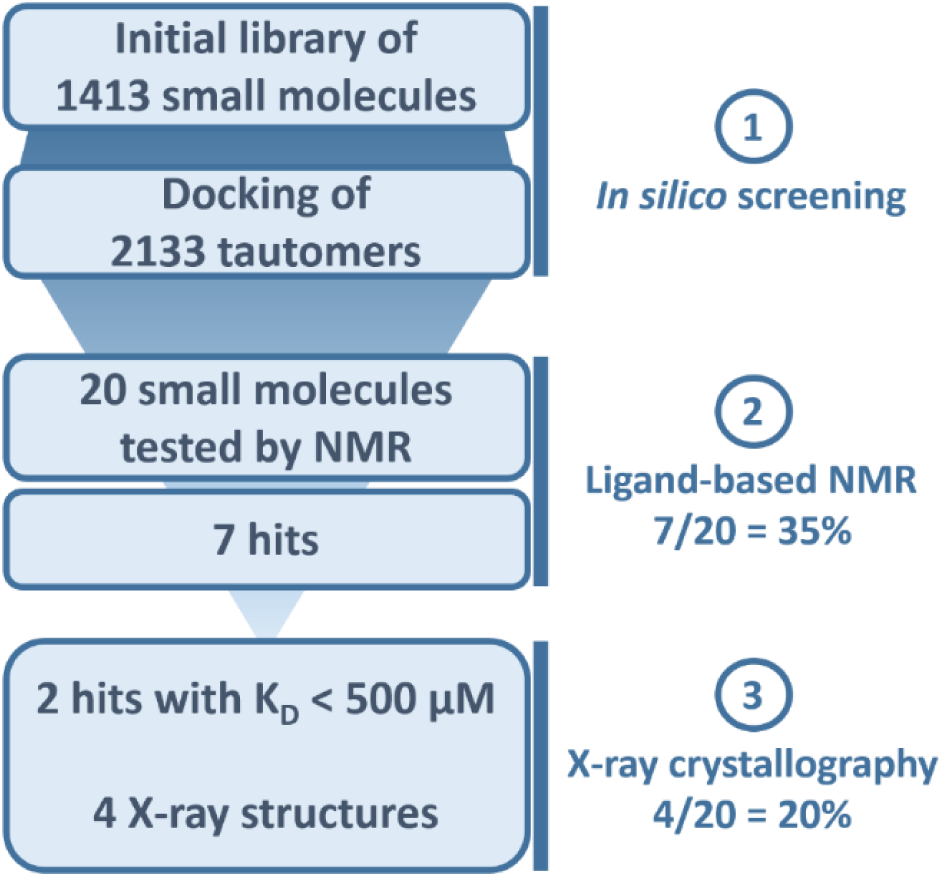
Flowchart of the combined *in silico* screening and experimental validation which consists of (1) high-throughput docking by the program SEED, ^9, 10^ (2) ligand-based NMR spectroscopy, and (3) final validation by a competition binding assay *in vitro*^11, 12^ and X-ray crystallography. The success ratio of the *in silico* screening is 35% according to ligand-observed NMR spectroscopy and 20% according to X-ray crystallography.

### *In vitro* Validation by Ligand-based NMR Spectroscopy and Measurements of Binding Affinity

The 20 small molecules were split into two mixtures at ten fragments each with minimal ^1^H NMR spectral overlap. These mixtures were subjected to ^1^H, STD (saturation transfer difference), and R_2_–filtered NMR experiments to determine fragment binding to BAZ2A.^13,14^ Compounds were classified as validated hits if at least one of these methods showed clear evidence of binding. This evaluation included competition experiments with **21** (GSK2801, Figure 2)^8^ to demonstrate specificity for the primary binding site.

**Figure 2.**
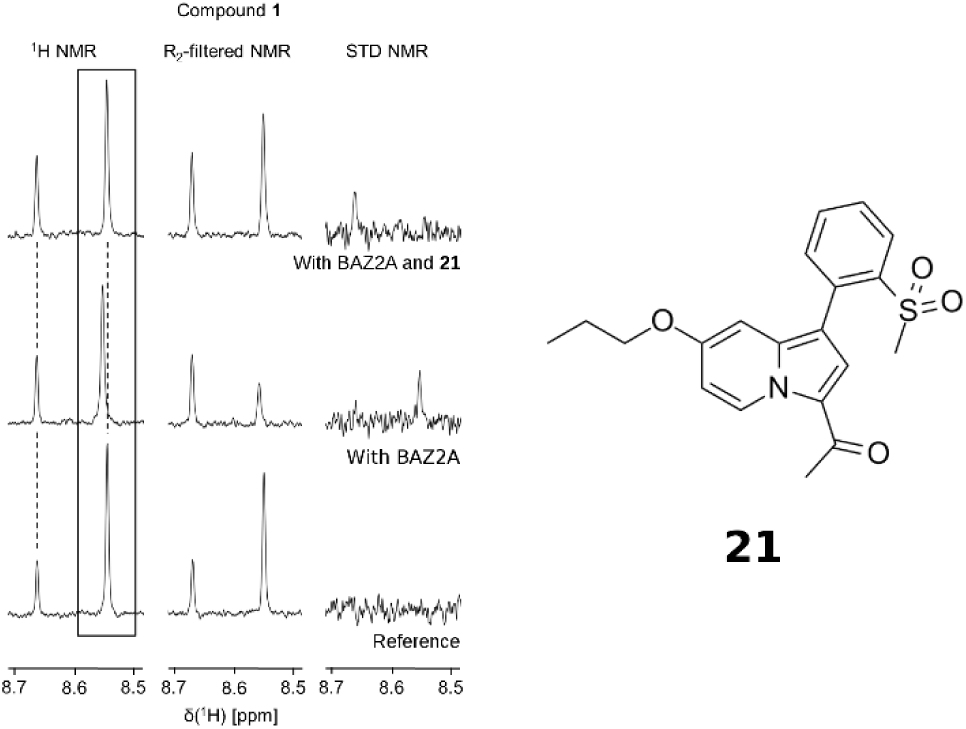
Ligand–observed NMR screening experiments for compound **1** which displays perturbation of the ^1^H chemical shift (left), an increased R2 relaxation (middle), and STD effects (right) in presence of BAZ2A. Titration of the nanomolar inhibitor **21** results in a reversal for all three observables, indicating specific binding in the Kac binding site. The depicted resonance at 8.55 ppm corresponds to H2 of the fragment hit **1**.

Overall, specific interactions with BAZ2A were observed for seven of the 20 fragments with R_2_-filtered NMR being the most sensitive technique (Table S2). The most pronounced effects were observed for compounds **1** and **2** showing STD effects and chemical shift perturbations (CSPs), as well as changes in the R_2_ relaxation (Table S2, Figure 2). The seven hits validated via NMR were further evaluated in an orthogonal assay *in vitro*. In this competitive binding assay based on DNA–tagged BAZ2A bromodomain and PCR quantification,^11, 12^ all compounds analyzed displayed significant competition at a concentration of 0.5 mM (Table S2). For the strongest competitors, compounds **1** and **2**, dissociation constants (KD) of 51 μM and 100 μM were measured, respectively (Table 1). Generally, affinity ranking via NMR is not straightforward for either CSPs, STD effects or changes in R_2_ relaxation when data from only a single ligand or receptor concentration is available. Yet, the pronounced effects for all three parameters are well in line with the K_D_ values measured for compounds **1** and **2**.

In the *in vitro* competition binding assay, similar KD values were measured for binding to the BAZ2A or the sequence-related BAZ2B bromodomain (Table 1, Figure S3). These results were to be expected because the bromodomains of BAZ2A and BAZ2B have very similar structure in the region of the Kac binding site that is occupied by compounds **1** and **2** (see below).

### X-ray Crystallography: Comparison with Poses Predicted by Docking

The structures of the bromodomain of BAZ2A in the complexes with compounds **1**–**4** were solved at 2.1 Å, 2.3 Å, 2.65 Å, and 2.8 Å resolution, respectively (Table S3, Figure S4). As predicted by docking, the four ligands occupy the Kac binding pocket with their aromatic ring sandwiched between the hydrophobic side chains of the so-called gatekeeper residue (Val1879) on one side of the pocket, and Val 1822 and Val 1827 on the other side (Figure 3). A carbonyl group in each of these fragment hits is involved in a water-bridged and direct hydrogen bond to the side chains of the conserved Tyr1830 and Asn1873, respectively. In addition, each of these compounds forms specific interactions with the Kac pocket.

**Figure 3.**
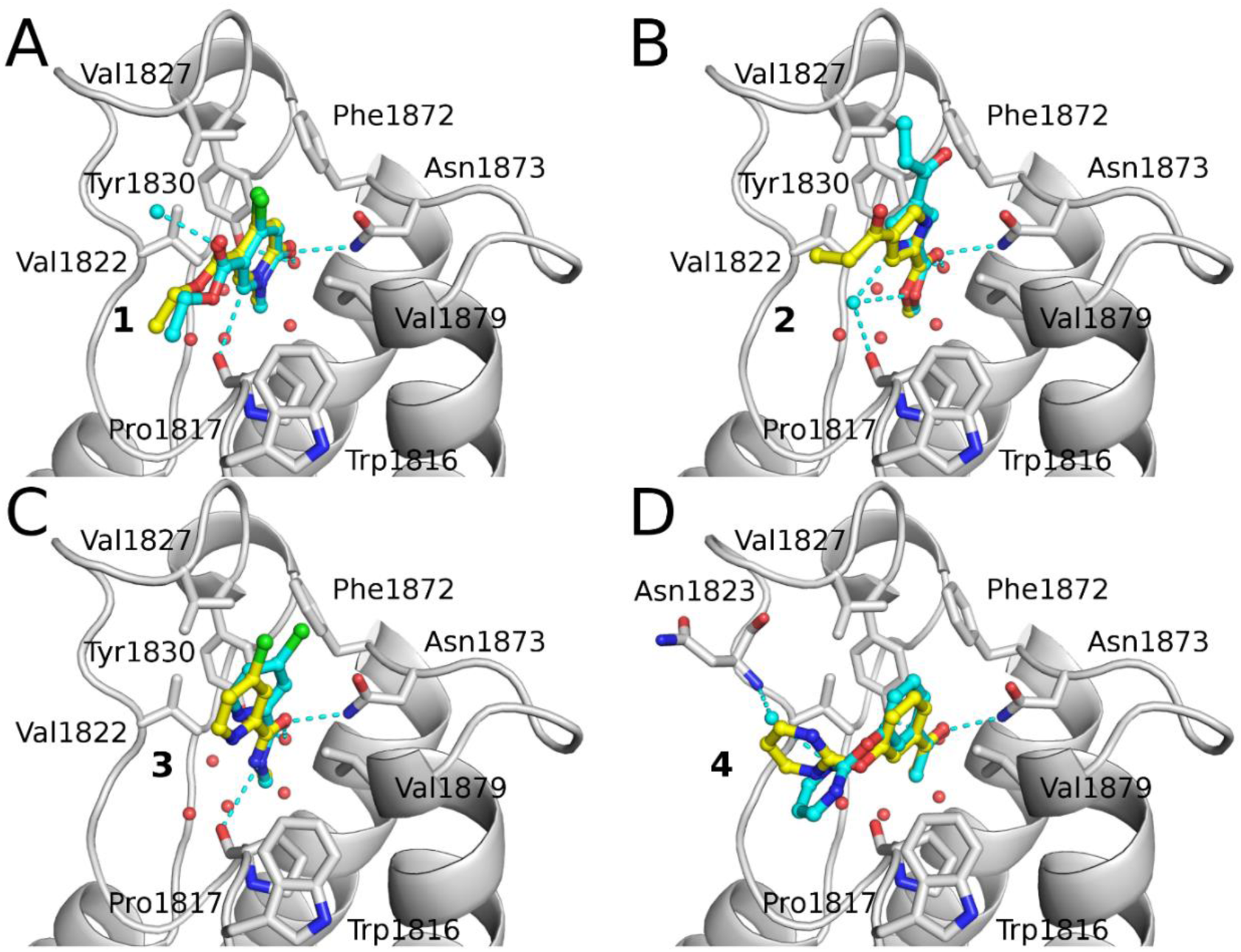
Structural validation of the fragment-based *in silico* screening campaign for BAZ2A. Comparison of the binding modes in the crystal structures (carbon atoms of ligands in cyan) with the binding poses predicted by docking with SEED,^9,10^ (carbon atoms in yellow) for compounds (A) **1**, (B) **2**, (C) **3**, and (D) **4** (PDB codes: 5MGJ, 5MGK, 5MGL, and 5MGM, respectively). Conserved water molecules (red spheres) and water molecules present in the crystal structure but not used for docking (cyan spheres) are shown. Compounds **2** and **4** have a different relative orientation of the two substituents, which could not be predicted by SEED since the compounds were docked as rigid molecules.

The 1-methylpyridinone derivative **1** has its 4-chloro and 3-ethyl-ester substituents pointing in part towards the solvent, but also in van der Waals contact with the side chains of Val1827 and Trp 1816–Pro1817, respectively (Figure 3A). The aromatic CH in position 2 is involved in a weak polar interaction with the carbonyl oxygen of Pro1817 (distance of 3.3 Å between the aromatic carbon of the ligand and the carbonyl oxygen). The pose predicted by docking is essentially identical to the binding mode observed in tthe crystal structure.

The 4-propionyl-1H-pyrrole derivative **2** is involved in hydrogen bonds with the conserved Tyr1830 and Asn1873 side chains through the carbonyl of the 2-methyl-ester substituent, while the propionyl group is directed towards the solvent (Figure 3B). The pyrrole nitrogen forms a water-mediated hydrogen bond with the main chain oxygen of Pro1817, and the same bridging water molecule is also in polar contact with the methoxy oxygen of the 2-methyl-ester substituent. The electron density is compatible also with the flipped binding mode (not shown), i.e., with the propionyl group buried and its carbonyl oxygen involved in the conserved hydrogen bonds. However, comparison of the two refined structures among them and with the refinement having both binding modes partially populated indicates that the binding mode with the deeply buried 2-methyl-ester substituent is preferred. Derivatives of acylpyrroles have already been reported as inhibitors of the bromodomain of the CREB-binding protein^15^ and BET bromodomains.^16, 17^ The binding mode of the previously reported acylpyrroles derivatives into their target is different with respect to the binding mode of compound **2** into the BAZ2A bromodomain, which is likely due to the different substituents and binding site side chains (Figure 4A, Figure S5A).

**Figure 4.**
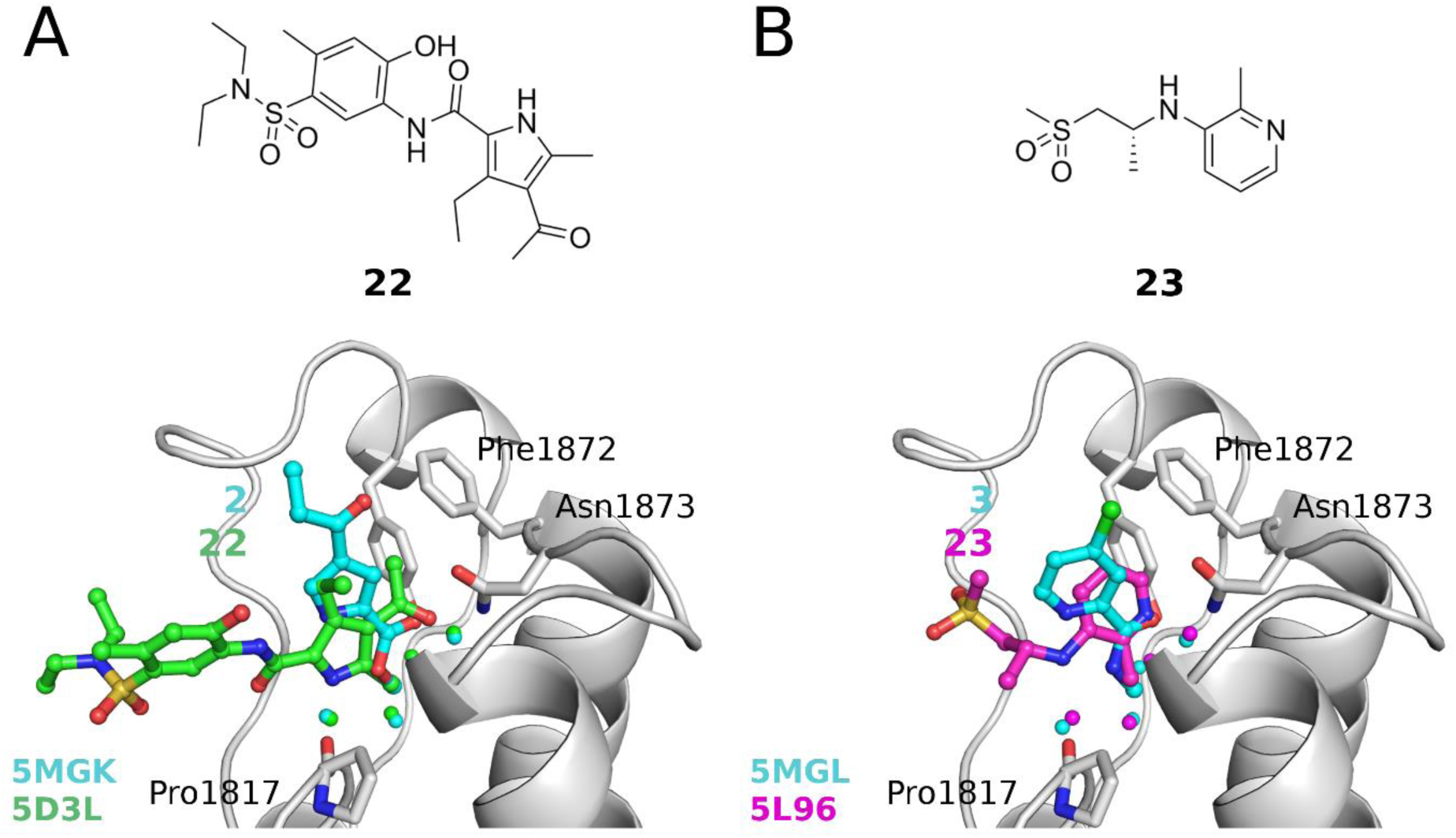
Comparison of the binding modes of the BAZ2A ligands **2** and **3** (carbon atoms in cyan) with previously reported bromodomain inhibitors **22** and **23**, respectively. (A) The overlap of the binding mode of compound **2** into the BAZ2A bromodomain (gray) with a representative of the tetrasubstituted pyrrole-series of BET bromodomain inhibitors (**22**, carbon atoms in green, 2D structure in the top panel)^17^ highlights two distinct binding modes. (B) The binding pose of the pyridine derivative **3** into the BAZ2A bromodomain (gray) is different with respect to the binding pose into BAZ2B of a 3-amino-2-methylpyridine bromodomain ligand (**23**, carbon atoms in magenta, 2D structure in the top panel). ^18^ In this figure and the following ones, the four-character labels are the PDB codes, and their colors correspond to those of the carbon atoms of the ligands. Structural overlap made use of the bromodomain Cα atoms, and only the BAZ2A bromodomain is shown to avoid overcrowding.

The pyridine derivative **3** (Figure 3C) forms a weak hydrogen bond with the carbonyl oxygen of Pro1817 (distance of 3.4 Å between the amide nitrogen in 3 and the carbonyl oxygen). Its 4-chloro substituent points towards the side chain of the conserved Phe1872 in the BC loop and in part towards the solvent. Because of the different substitution patterns, the binding mode of the pyridine derivative **3** in BAZ2A is completely different than the recently disclosed complexes of BAZ2B with 3-amino-2-methylpyridine derivatives (Figure 4B, Figure S5B).^18^ The compound **3** has its carbonyl oxygen at hydrogen-bond distance to the side chains of the conserved Tyr1830 (water-bridged) and Asn1873 of BAZ2A while for the 3-amino-2-methylpyridine derivatives (which are devoid of carbonyl substituents) it is the pyridine nitrogen that is involved in the analogous hydrogen bonds with the conserved Tyr1901 (water-bridged) and Asn1944 of BAZ2B.

The acetylbenzene derivative **4** (Figure 3D) has its 3-oxypyrimidine substituent pointing towards the side chains of Trp1816 and Pro1817. One of the pyrimidine nitrogen is involved in a water-bridged hydrogen bond with the backbone NH of Asn1823. A similar water-bridged hydrogen bond has been reported in crystal structures of BAZ2B/ligand complexes (Figure S6).^6,8,18^

### X-ray Crystallography: Comparison of binding modes in BAZ2A and BAZ2B

Structures of compounds **1**, **2**, and **3** in complex with BAZ2B were also determined at 1.95 Å, 1.9 Å, and 2.1 Å resolution, respectively (Table S3). The comparison of the BAZ2A and BAZ2B crystal structures provides further evidence of the influence of the gatekeeper side chain on the orientation of small-molecule ligands. These two homologous bromodomains have almost identical structure in the acetyl-lysine binding site. The most relevant difference is the gatekeeper residue, which is Val1879 in BAZ2A and Ile1950 in BAZ2B. In agreement with previous crystallographic evidence,^19^ the additional methyl group of the latter side chain forces a tilting of the aromatic rings of the fragment hits (20° in compounds **1** and **2**, and 10° in compound **3**) which is coupled to a slight displacement of the ZA loop (Figure 5). The largest deviation in the backbone is observed for the BAZ2B residue Lys1896 in the C-terminal segment of the ZA loop, whose Cα position deviates from the corresponding residues in BAZ2A (Arg1825) by 1.8 Å, 1.6 Å, and 1.6 Å in the structures with compounds **1**, **2**, and **3**, respectively (Figure 5A-D).

**Figure 5.**
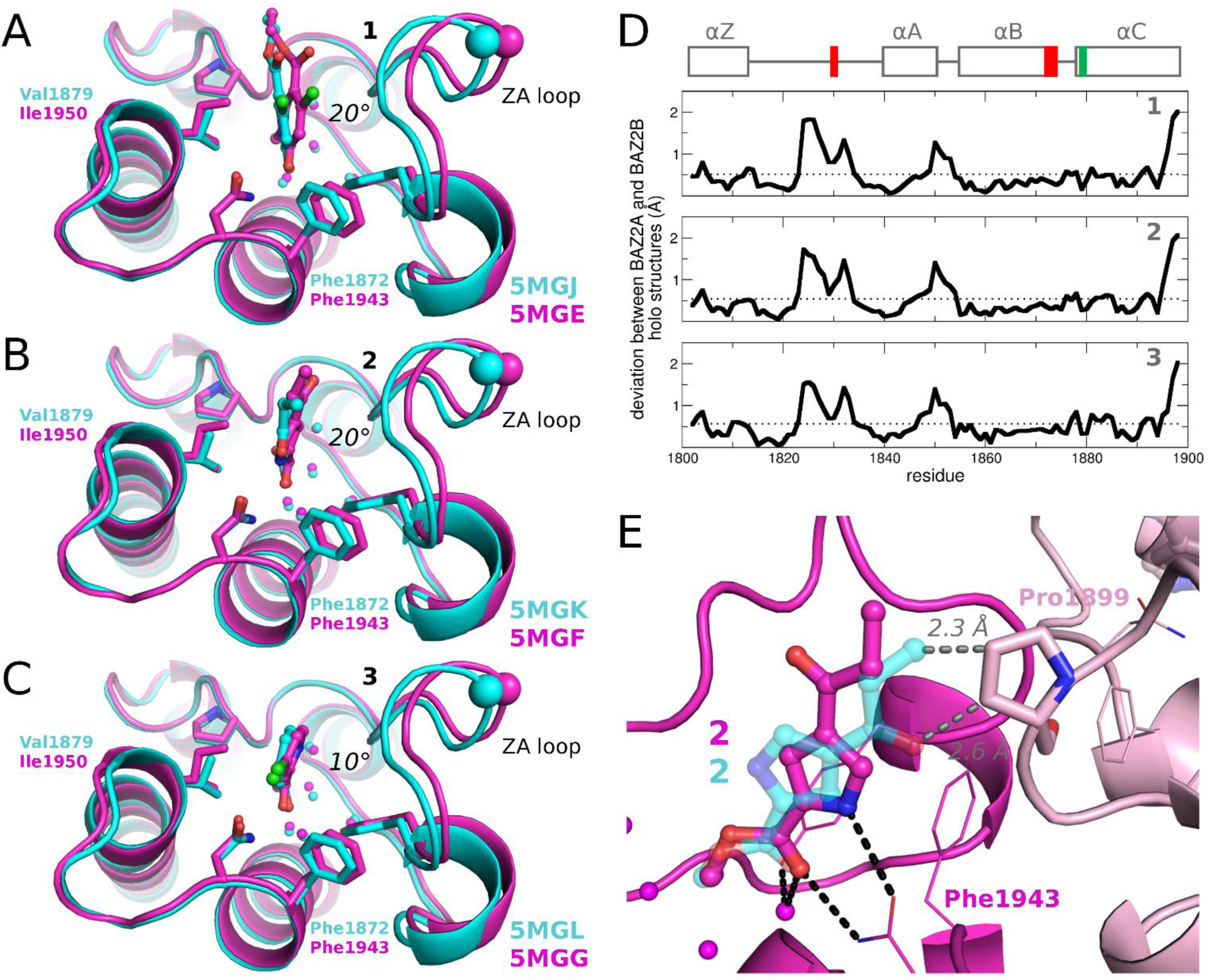
(A-C) Comparison of the binding modes of compounds (A) **1**, (B) **2**, and (C) **3** in BAZ2A (cyan) and BAZ2B (magenta). The gatekeeper residue is Val1879 in BAZ2A and Ile 1950 in BAZ2B. The additional methyl group in the latter side chain influences the orientation of the ligand, position of the ZA loop, and orientation of the conserved Phe in the BC loop, *viz*., Phe1872 and Phe1943 in BAZ2A and BAZ2B, respectively. The Cα atoms of BAZ2A Arg1825 and BAZ2B Lys1896 (cyan and magenta sphere, respectively) are shown to emphasize the segment of the ZA loop with the largest displacement. (D) Profile of the Cα atoms deviation between BAZ2A and BAZ2B for the complexes with compounds **1** (top), **2** (middle), and **3** (bottom). The helical segments are shown (rectangles in the top) together with the location along the sequence of the highly conserved side chains and the gatekeeper residue (red and green, respectively). The numbering in the x-axis refers to the BAZ2A bromodomain, and for each system the average displacement is also shown (dotted line). (E) The binding mode of compound **2** in BAZ2A (carbon atoms in cyan) is not compatible with the crystallographic packing of BAZ2B (magenta): the fragment hit **2** would clash with the side chain of Pro1899 of a neighboring bromodomain molecule (pink). Interatomic distances are shown with grey dashed lines. In all panels the bromodomain structures were superimposed using the Cα atoms of the residues in α–helical conformation.

In BAZ2B, the fragment hit **2** has its pyrrole nitrogen in hydrogen bond contact with the side chain oxygen of the conserved Asn1944 while the pyrrole is oriented differently in BAZ2A. It has however to be noticed that the binding mode observed in BAZ2A is not compatible with the crystallographic packing of BAZ2B, where the propionyl group would clash with a neighboring protein chain (Figure 5E). It remains then questionable whether the different orientation of the pyrrole ring in the complexes with BAZ2A and BAZ2B has to be ascribed to the difference in the gatekeeper residue or to the crystallographic constraints.

### Suggestions for Fragment Growing

Structural overlap of the BAZ2A and BAZ2B bromodomains shows that besides the aforementioned difference in the gatekeeper residue, there are only two additional differences in the Kac binding site, both of which are in the ZA loop: Glu1820/Leu1891 and Ser1828/Pro1899 (in BAZ2A/BAZ2B, respectively). The binding modes of fragment hits **1** and **3** suggest optimization of selectivity by growing these hits into the direction of Glu1820/Leu1891 and/or Ser1828/Pro1899 (Figure 6). In particular, amide coupling in position 3 of the 1-methylpyridinone derivative **1** could be exploited to position a positively charged amino group or hydrophobic moiety in proximity of Glu1820 (BAZ2A) or Leu1891 (BAZ2B), respectively (Figure 6A). The ionic interaction with Glu1820 would require reorientation of its side chain which should be possible because it is solvent-exposed and not involved in intra-protein interactions. Concerning the Ser1828/Pro1899 difference, both pyridine-based hits **1** and **3** could be elongated along the direction of their 4-chlorine substituent (green atom in Figure 6) which could result in a hydrogen bond or hydrophobic contact with the side chains of Ser1828 (BAZ2A) or Pro1899 (BAZ2B), respectively.

**Figure 6.**
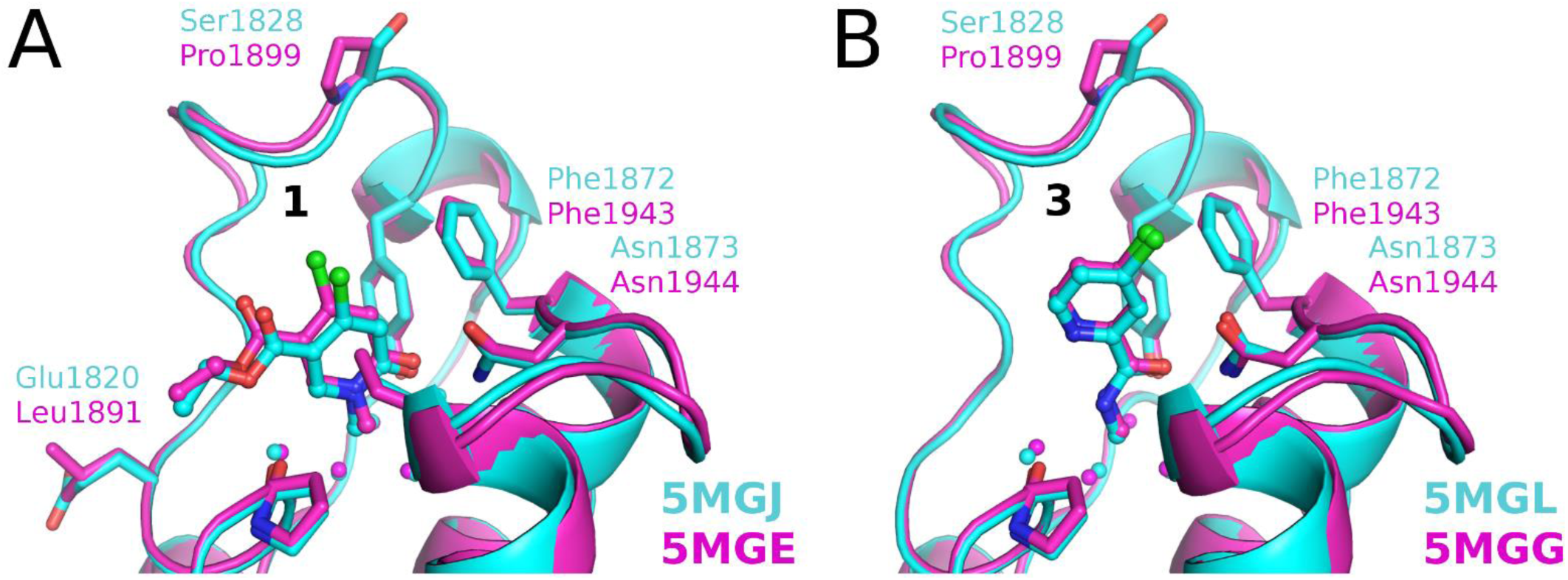
Comparison of the crystal structures of ligands **1** (A) and **3** (B) complexed to BAZ2A (cyan) and BAZ2B (magenta) suggests fragment growing to achieve binding selectivity. Future efforts aimed to achieve selective ligands for the BAZ2A or BAZ2B bromodomains could direct the synthesis of derivatives of fragment hits to form polar interactions with the side chains of Glu1820 and/or Ser1828 (BAZ2A) or hydrophobic contacts with Leu1891 and/or Pro1899 (BAZ2B).

## CONCLUSIONS

We have used a combined computational and experimental strategy to identify small molecules that bind to the bromodomain of BAZ2A. Starting from a library of nearly 1500 small molecules, 20 compounds were selected by high-throughput docking as potential Kac-competitive binders, and seven of them were confirmed by ligand-based NMR spectroscopy. Furthermore, the compounds **1** and **2** have a ligand efficiency of 0.42 kcal/mol per non-hydrogen atom for the BAZ2A bromodomain in a competition binding assay *in vitro*. For the ligands **1**–**4** we could solve the crystal structure in the complex with the BAZ2A bromodomain which confirmed the predicted binding modes. Thus, the *in silico* screening had a success ratio of 10% if one considers the affinity as measured by the *in vitro* assay, 20% according to the evidence of binding to the target as provided by the crystal structure of the complex, or 35% according to ligand-observed NMR spectroscopy. It is important to note that the computational screening is very efficient as the automatic docking of the 1413 compounds required only two hours on a single core of a commodity computer.

We also solved the crystal structures of the complex of ligands **1**, **2**, and **3** with the BAZ2B bromodomain which differs from BAZ2A mainly in the gatekeeper residue (Val1879 and Ile1950 in BAZ2A and BAZ2B, respectively). The comparison of the binding mode in BAZ2A and BAZ2B provides further evidence of the influence of the gatekeeper side chain on the orientation of the head group (Figure 5).^19^

To the best of our knowledge, the structures of the complexes of the BAZ2A bromodomain with ligands **1**–**4** are the first *holo* structures of this bromodomain except for the complex of BAZ2A with a diacetylated histone 4 peptide (H4_14–22_K16acK20ac) (PDB code 4QBM). The only interactions shared by the histone peptide and our ligands are those of the acetylated side chain of Lys16 which occupies the Kac binding site while the rest of the peptide is bound to the surface and oriented towards the BC loop (Figure S7). Thus, our crystal structures provide further, novel information on hydrophobic contacts and (water-bridged) hydrogen bonds.

Our fragment hits show similar affinity for the BAZ2A and BAZ2B bromodomains which have very similar Kac binding site. Interestingly, the crystal structures of the complexes with the 1-methylpyridinone derivative **1** and 4-chloropyridine derivative **3** suggest fragment growing into either or both of two directions to reach the two residues in the Kac binding site that differ between BAZ2A and BAZ2B (Figure 6). These suggestions for hit expansion are expected to provide selectivity between two bromodomains in the same sub-family.

## EXPERIMENTAL SECTION

### Fragment Docking and Selection

A library of 1413 small molecules was considered for in silico screening (distributions of properties, *e. g.*, molecular weight, are shown in Figure S1). These molecules originate from an existing set of fragments maintained by the laboratory of one of the authors (C. R.). The library was selected for chemical diversity from a large panel of commercial suppliers and academic collaborators. Quality controls ensuring identity, stability and solubility were performed. Next, a total of 2133 tautomers were generated using the calculator plugin of Marvin (Figure 1).^20^ The structure of the BAZ2A bromodomain used for docking is the one in the complex with a diacetylated histone H4_14–22_K16acK20ac peptide (PDB code: 4QBM).^21^ The binding site definition for SEED docking consisted of the conserved asparagine in the BC loop (Asn1873) and the five conserved water molecules in the Kac binding site. The partial charges and van der Waals parameters for the atoms in the protein and the small molecules were taken from the CHARMM36 all-atom force field^22, 23^ and the CHARMM general force field (CGenFF),^24^ respectively. Importantly, the CHARMM36 force field and CGenFF are fully consistent in their partial charges and van der Waals parameters. The evaluation of the binding energy in the program SEED^9, 10^ consists of a force field-based energy function with a continuum dielectric approximation of desolvation penalties by the generalized Born model.^25^ The values of the dielectric constant were 2.0 and 78.5 for the volume occupied by the solute and solvent, respectively. The docking of 2133 tautomers of 1413 small molecules with SEED required approximately 2 h (about 3 s per compound) of a single core of a Xeon E5410 processor at 2.33 GHz.

Compounds were ranked using a consensus scoring function based on the median of the ranks of three terms calculated by SEED, namely (i) the total binding energy, (ii) the intermolecular van der Waals contribution divided by the number of non-hydrogen atoms of the ligand (van der Waals efficiency), and (iii) the difference between the electrostatic contribution to the intermolecular interaction energy in the solvent and the solvation energy of the ligand. We used the median value of the rankings as it is less sensitive to outliers than the mean value and has shown robustness in previous *in silico* high-throughput docking campaigns.^26,27^

### Compound Purity

All compounds were purchased form Key Organics. The purity of the molecules was analyzed by HPLC-MS and is ≥ 95%. NMR spectra were analyzed for the presence of contaminants.

### Protein Expression and Purification

BAZ2AA–c002 was a gift from Nicola Burgess-Brown (Addgene plasmid #53623). For crystallization purposes, BAZ2B bromodomain (aa. 1858–1972) was produced as detailed in ^28^. For BAZ2A bromodomain (aa. 1796–1899), after induction with IPTG 0.2 mM and overnight expression at 20°C, the 6×His –tagged protein was purified by IMAC and eluted with a linear gradient of imidazole. Buffer was exchanged and the 6×His tag removed with tobacco etch virus protease. The bromodomain was further purified by a second IMAC and a size-exclusion chromatography using a Superdex 75 column. Proteins was concentrated to 23 mg/mL and frozen in liquid nitrogen.

### NMR Spectroscopy

NMR spectra were recorded on a PremiumCompact 600 MHz spectrometer equipped with a OneNMR probe (Agilent) and spectra were processed using the MestReNova software suite from Mestrelab Research S. L.^29^ All NMR experiments were conducted at 25°C. A DPFGSE pulse sequence was utilized for solvent suppression. ^30^ STD NMR experiments utilized a train of 50 ms Gauss pulses for a saturation time t_sat_ of 4.0 s.^13^ The on-resonance irradiation frequency ν_sat_ was set to 0.0 ppm, while the off-resonance irradiation frequency ν_ref_ was set to 80.0 ppm. The relaxation delay d1 was set to 0.0 s and the acquisition time t_acq_ was set to 2.0 s. Receptor resonances were suppressed via a T_1,rho_ filter of 35 ms duration. For each spectrum 256 scans were recorded in 5 mm sample tubes at sample volumes of 500 to 550 μL. R_2_–filtered NMR experiments were implemented utilizing the Carr–Purcell–Meiboom–Gill (CPMG) pulse sequence.^31^ The relaxation delay d1 was set to 2.0 s and the acquisition time t_acq_ was set to 2.0 s. Spectra were recorded at 128 scans at a frequency of 180° ν_CPMG_ of 100 Hz and a relaxation time T of 0.4 s under the same sample conditions as indicated for the STD NMR spectra.

The induction of CSPs in presence of BAZ2A was analyzed using regular ^1^H NMR experiments. The relaxation delay d_1_ was set to 2.0 s and the acquisition time t_acq_ was set to 2.0 s. Spectra were recorded at 128 scans under the same sample conditions as indicated for the STD NMR spectra.

A genetic algorithm was utilized to generate two fragment mixtures with minimal ^1^H NMR spectral overlap from the 20 selected compounds. These mixtures were prepared at 200 μM fragment concentration in 50 mM H_3_PO_4_ with 100% D_2_O, 2% DMSO–d_6_ and 150 mM NaCl at pH 7.4. TSP–d_4_ served as an internal reference at a concentration of 100 μM. Resonances of the analyzed fragment mixtures were assigned by comparison to previously acquired H^1^ NMR spectra of the individual fragments. The stability of the fragment mixtures over 16 h at room temperature was monitored via ^1^H NMR. Next, R_2_–filtered and STD NMR experiments for the fragment mixtures were conducted as described above, first in absence of BAZ2A, then repeated after the addition of 20 μM BAZ2A and finally in a consecutive experiment in presence of 40 μM of the known nanomolar ligand **21**.^8^

Compounds were defined as validated hits, if either an STD effect, an increased R_2,obs_ value or a CSP was observed in presence of BAZ2A, and if these effects were reversed by addition of compound **21**. Moreover, only fragments that did not display an STD effect in absence of BAZ2A were selected.

### BROMOscan Assay

BROMOscan is a competition-based technology using a ligand immobilized to a solid support and DNA-tagged bromodomains. Bromodomains are incubated with the ligand in the presence and absence of the putative inhibitors and eluted to be quantified by qPCR. Small molecules inhibiting the bromodomain binding to the immobilized ligand will reduce the amount of bromodomain captured and thus the qPCR signal.^11, 12^ Dissociation constants (K_D_) were calculated by fitting a 12-point dilution curve with starting concentration of 0.5 mM and dilution factor of 3.0. All dose-responses were measured in duplicates.

### Crystallization, Data Collection and Structure Solution

Crystallization and soaking for the BAZ2B bromodomain were performed as previously described.^18^ For the BAZ2A bromodomain, showers of extremely thin needles were obtained at 4°C in Tris pH 8, MgCl_2_ 0.2 M, PEG3350 26%. These were subsequently used for microseeding, obtaining more disperse and slightly larger needles in Tris pH 7.5, MgCl_2_ 0.2 M, PEG3350 18–22%. Complexes with the compounds of interest (50 mM or saturating solutions for less soluble compounds) were obtained by co-crystallization. Compounds were dissolved in the crystallization solution devoid of DMSO, which does bind in the Kac pocket of bromodomains.^32^ Co-crystals were cryoprotected with ethylene glycol and frozen in liquid nitrogen.

Diffraction data were collected at the Elettra Synchrotron Light Source (Trieste, Italy), XRD1 beamline. Data were processed with either XDS^33^ or MOSFLM,^34^ and Aimless;^35^ high resolution cutoff was selected according to Karplus and Diederichs.^36, 37^ Structures were solved by molecular replacement with Phaser^38^ using PDB 4IR5 as search model for BAZ2B and PDB 4LZ2 for BAZ2A. Initial models were refined alternating cycles of automatic refinement with either Phenix^39^ or REFMAC^40^ and manual model building with COOT.^41^

## ASSOCIATED CONTENT

### Supporting Information

The Supporting Information contains:

Biophysical evaluation of the seven compounds selected by the high-throughput docking, X-ray crystal structure refinement data, distribution of the chemical properties of the libraries, insights into the energetic contributions calculated *in silico*, and structural analyses of all reported crystal structures (PDF). Molecular formula strings for ligands **1**–**20** (CSV).

### Accession Codes

PDB ID codes: 5MGJ (BAZ2A-1), 5MGK (BAZ2A-2), 5MGL (BAZ2A-3), 5MGM (BAZ2A-4), 5MGE (BAZ2B-1), 5MGF (BAZ2B-2), and 5MGG (BAZ2B3).

## Notes

The authors declare no competing financial interest.

## ACKNOWLEDGMENTS

We thank Jonas Aretz, Nicholas Deerain, and Jean-Rémy Marchand for technical support and interesting discussions. We thank the Structural Genomics Consortium at University of Oxford for providing the plasmid of the BAZ2B bromodomain. Diffraction data were acquired on the XRD1 beamline, ELETTRA Synchrotron Light Source in Trieste, Italy. This work was supported financially by a grant of the Swiss National Science Foundation to A.C. (grant 31003A_169007). D.S. is a recipient of the Systems X.ch translational postdoc fellowship and gratefully acknowledges support from the Holcim Foundation. C.R. and E.W. thank the Max Planck Society and the German Research Foundation (DFG, RA1944/2–1) for financial support.

## ABBREVIATIONS

BAZ2A: bromodomain adjacent to zinc finger domain protein 2A
BAZ2B: bromodomain adjacent to zinc finger domain 2B
BET: bromodomain and extra terminal
CPMG: Carr-Purcell-Meiboom-Gill sequence
CREB: cAMP response element-binding protein
CSP: chemical shift perturbation
DMSO: dimethyl sulfoxide
DPFGSE: Double Pulsaed Field Gradient Spin Echo sequence
IMAC: immobilized metal affinity chromatography
IPTG: isopropyl β-D-1-thiogalactopyranoside
Kac: acetyllysine
LE: ligand efficiency
STD: saturation transfer difference
TSP: trimethylsilylpropanoic acid.

## REFERENCES

1. Dhalluin, C.; Carlson, J. E.; Zeng, L.; He, C.; Aggarwal, A. K.; Zhou, M. M. Structure and ligand of a histone acetyltransferase bromodomain. Nature 1999, 399, 491–496.

2. Filippakopoulos, P.; Picaud, S.; Mangos, M.; Keates, T.; Lambert, J. P.; Barsyte-Lovejoy, D.; Felletar, I.; Volkmer, R.; Muller, S.; Pawson, T.; Gingras, A. C.; Arrowsmith, C. H.; Knapp, S. Histone recognition and large-scale structural analysis of the human bromodomain family. Cell 2012, 149, 214–231.

3. Flynn, E. M.; Huang, O. W.; Poy, F.; Oppikofer, M.; Bellon, S. F.; Tang, Y.; Cochran, A. G. A subset of human bromodomains recognizes butyryllysine and crotonyllysine histone peptide modifications. Structure 2015, 23, 1801–1814.

4. Gu, L.; Frommel, S. C.; Oakes, C. C.; Simon, R.; Grupp, K.; Gerig, C. Y.; Bar, D.; Robinson, M. D.; Baer, C.; Weiss, M.; Gu, Z.; Schapira, M.; Kuner, R.; Sultmann, H.; Provenzano, M.; Cancer, I. P. o. E. O. P.; Yaspo, M. L.; Brors, B.; Korbel, J.; Schlomm, T.; Sauter, G.; Eils, R.; Plass, C.; Santoro, R. BAZ2A (TIP5) is involved in epigenetic alterations in prostate cancer and its overexpression predicts disease recurrence. Nat. Genet. 2015, 47, 22–30.

5. Ferguson, F. M.; Fedorov, O.; Chaikuad, A.; Philpott, M.; Muniz, J. R. C.; Felletar, I.; von Delft, F.; Heightman, T.; Knapp, S.; Abell, C.; Ciulli, A. Targeting low druggability bromodomains: fragment based screening and inhibitor design against the BAZ2B bromodomain. J. Med. Chem. 2013, 56, 10183–10187.

6. Drouin, L.; McGrath, S.; Vidler, L. R.; Chaikuad, A.; Monteiro, O.; Tallant, C.; Philpott, M.; Rogers, C.; Fedorov, O.; Liu, M.; Akhtar, W.; Hayes, A.; Raynaud, F.; Muller, S.; Knapp, S.; Hoelder, S. Structure enabled design of BAZ2-ICR, a chemical probe targeting the bromodomains of BAZ2A and BAZ2B. J. Med. Chem. 2015, 58, 2553–2559.

7. Chung, C.-w.; Dean, A. W.; Woolven, J. M.; Bamborough, P. Fragment-based discovery of bromodomain inhibitors part 1: inhibitor binding modes and implications for lead discovery. J. Med. Chem. 2011, 55, 576–586.

8. Chen, P.; Chaikuad, A.; Bamborough, P.; Bantscheff, M.; Bountra, C.; Chung, C. W.; Fedorov, O.; Grandi, P.; Jung, D.; Lesniak, R.; Lindon, M.; Muller, S.; Philpott, M.; Prinjha, R.; Rogers, C.; Selenski, C.; Tallant, C.; Werner, T.; Willson, T. M.; Knapp, S.; Drewry, D. H. Discovery and characterization of GSK2801, a selective chemical probe for the bromodomains BAZ2A and BAZ2B. J. Med. Chem. 2016, 59, 1410–1424.

9. Majeux, N.; Scarsi, M.; Apostolakis, J.; Ehrhardt, C.; Caflisch, A. Exhaustive docking of molecular fragments with electrostatic solvation. Proteins: Struct., Funct., Genet. 1999, 37, 88–105.

10. Majeux, N.; Scarsi, M.; Caflisch, A. Efficient electrostatic solvation model for protein-fragment docking. Proteins: Struct., Funct., Genet. 2001, 42, 256–268.

11. Fabian, M. A.; Biggs, W. H., 3rd; Treiber, D. K.; Atteridge, C. E.; Azimioara, M. D.; Benedetti, M. G.; Carter, T. A.; Ciceri, P.; Edeen, P. T.; Floyd, M.; Ford, J. M.; Galvin, M.; Gerlach, J. L.; Grotzfeld, R. M.; Herrgard, S.; Insko, D. E.; Insko, M. A.; Lai, A. G.; Lelias, J. M.; Mehta, S. A.; Milanov, Z. V.; Velasco, A. M.; Wodicka, L. M.; Patel, H. K.; Zarrinkar, P. P.; Lockhart, D. J. A small molecule-kinase interaction map for clinical kinase inhibitors. Nat. Biotechnol. 2005, 23, 329–336.

12. Quinn, E.; Wodicka, L.; Ciceri, P.; Pallares, G.; Pickle, E.; Torrey, A.; Hunt, J.; Treiber, D. BROMOscan - a high throughput, quantitative ligand binding platform identifies best-in-class bromodomain inhibitors from a screen of mature compounds targeting other protein classes. Cancer Res. 2013, 73, 4238.

13. Mayer, M.; Meyer, B. Characterization of ligand binding by saturation transfer difference NMR spectroscopy. Angew. Chem., Int. Ed. 1999, 38, 1784–1788.

14. Hajduk, P. J.; Olejniczak, E. T.; Fesik, S. W. One-dimensional relaxation- and diffusion-edited NMR methods for screening compounds that bind to macromolecules. J. Am. Chem. Soc. 1997, 119, 12257–12261.

15. Xu, M.; Unzue, A.; Dong, J.; Spiliotopoulos, D.; Nevado, C.; Caflisch, A. Discovery of CREBBP bromodomain inhibitors by high-throughput docking and hit optimization guided by molecular dynamics. J. Med. Chem. 2016, 59, 1340–1349.

16. Lucas, X.; Wohlwend, D.; Hägle, M.; Schmidtkunz, K.; Gerhardt, S.; Schäle, R.; Jung, M.; Einsle, O.; Gänther, S. 4-Acyl pyrroles: mimicking acetylated lysines in histone code reading. Angew. Chem., Int. Ed. 2013, 52, 14055–14059.

17. Hägle, M.; Lucas, X.; Weitzel, G.; Ostrovskyi, D.; Breit, B.; Gerhardt, S.; Einsle, O.; Gunther, S.; Wohlwend, D. 4-Acyl pyrrole derivatives yield novel vectors for designing inhibitors of the acetyl-lysine recognition site of BRD4(1). J. Med. Chem. 2016, 59, 1518–1530.

18. Marchand, J. R.; Lolli, G.; Caflisch, A. Derivatives of 3-amino-2-methylpyridine as BAZ2B bromodomain ligands: in silico discovery and in crystallo validation. J. Med. Chem. 2016, 59, 9919–9927.

19. Unzue, A.; Zhao, H.; Lolli, G.; Dong, J.; Zhu, J.; Zechner, M.; Dolbois, A.; Caflisch, A.; Nevado, C. The “gatekeeper” residue influences the mode of binding of acetyl indoles to bromodomains. J. Med. Chem. 2016, 59, 3087–3097.

20. MarvinSketch 15.8.17, ChemAxon, Budapest, Hungary, 2015.

21. Tallant, C.; Valentini, E.; Fedorov, O.; Overvoorde, L.; Ferguson, F. M.; Filippakopoulos, P.; Svergun, D. I.; Knapp, S.; Ciulli, A. Molecular basis of histone tail recognition by human TIP5 PHD finger and bromodomain of the chromatin remodeling complex NoRC. Structure 2015, 23, 80–92.

22. MacKerell, A. D.; Bashford, D.; Bellott, M.; Dunbrack, R. L.; Evanseck, J. D.; Field, M. J.; Fischer, S.; Gao, J.; Guo, H.; Ha, S.; Joseph-McCarthy, D.; Kuchnir, L.; Kuczera, K.; Lau, F. T.; Mattos, C.; Michnick, S.; Ngo, T.; Nguyen, D. T.; Prodhom, B.; Reiher, W. E.; Roux, B.; Schlenkrich, M.; Smith, J. C.; Stote, R.; Straub, J.; Watanabe, M.; Wiorkiewicz-Kuczera, J.; Yin, D.; Karplus, M. All-atom empirical potential for molecular modeling and dynamics studies of proteins. J. Phys. Chem. B 1998, 102, 3586–3616.

23. MacKerell, A. D.; Feig, M.; Brooks, C. L., 3rd. Improved treatment of the protein backbone in empirical force fields. J. Am. Chem. Soc. 2004, 126, 698–699.

24. Vanommeslaeghe, K.; Hatcher, E.; Acharya, C.; Kundu, S.; Zhong, S.; Shim, J.; Darian, E.; Guvench, O.; Lopes, P.; Vorobyov, I.; Mackerell, A. D. CHARMM general force field: a force field for drug-like molecules compatible with the CHARMM all-atom additive biological force fields. J. Comput. Chem. 2010, 31, 671–690.

25. Scarsi, M.; Apostolakis, J.; Caflisch, A. Continuum electrostatic energies of macromolecules in aqueous solutions. J. Phys. Chem. A 1997, 101, 8098–8106.

26. Huang, D.; Caflisch, A. Library screening by fragment-based docking. J. Mol. Recognit. 2010, 23, 183–193.

27. Zhao, H.; Caflisch, A. Molecular dynamics in drug design. Eur. J. Med. Chem. 2015, 91, 4–14.

28. Lolli, G.; Caflisch, A. High-throughput fragment docking into the BAZ2B bromodomain: efficient in silico screening for X-ray crystallography. ACS Chem. Biol. 2016, 11, 800–807.

29. MestreNova 10.0.2, MestreLab Research, Santiago de Compostela, Spain, 2015.

30. Hwang, T. L.; Shaka, A. J. Water suppression that works-excitation sculpting using arbitrary wave-forms and pulsed-field gradients. J. Magn. Reson., Ser. A 1995, 112, 275–279.

31. Carr H. Y.; M., P. E. Effects of diffusion on free precession in nuclear magnetic resonance experiments. Phys. Rev. 1954, 94, 630–638.

32. Lolli, G.; Battistutta, R. Different orientations of lowmolecular-weight fragments in the binding pocket of a BRD4 bromodomain. Acta Crystallogr., Sect. D: Biol. Crystallogr. 2013, 69, 2161–2164.

33. Kabsch, W. Xds. Acta Crystallogr., Sect. D: Biol. Crystallogr. 2010, 66, 125–132.

34. Battye, T. G.; Kontogiannis, L.; Johnson, O.; Powell, H. R.; Leslie, A. G. iMOSFLM: a new graphical interface for diffraction-image processing with MOSFLM. Acta Crystallogr., Sect. D: Biol. Crystallogr. 2011, 67, 271–281.

35. Evans, P. R.; Murshudov, G. N. How good are my data and what is the resolution? Acta Crystallogr., Sect. D: Biol. Crystallogr. 2013, 69, 1204–1214.

36. Karplus, P. A.; Diederichs, K. Linking crystallographic model and data quality. Science 2012, 336, 1030–1033.

37. Karplus, P. A.; Diederichs, K. Assessing and maximizing data quality in macromolecular crystallography. Curr. Opin. Struct. Biol. 2015, 34, 60–68.

38. McCoy, A. J.; Grosse-Kunstleve, R. W.; Adams, P. D.; Winn, M. D.; Storoni, L. C.; Read, R. J. Phaser crystallographic software. J. Appl. Crystallogr. 2007, 40, 658–674.

39. Adams, P. D.; Afonine, P. V.; Bunkoczi, G.; Chen, V. B.; Davis, I. W.; Echols, N.; Headd, J. J.; Hung, L. W.; Kapral, G. J.; Grosse-Kunstleve, R. W.; McCoy, A. J.; Moriarty, N. W.; Oeffner, R.; Read, R. J.; Richardson, D. C.; Richardson, J. S.; Terwilliger, T. C.; Zwart, P. H. PHENIX: a comprehensive Python-based system for macromolecular structure solution. Acta Crystallogr., Sect. D: Biol. Crystallogr. 2010, 66, 213–221.

40. Murshudov, G. N.; Vagin, A. A.; Dodson, E. J. Refinement of macromolecular structures by the maximum-likelihood method. Acta Crystallogr., Sect. D: Biol. Crystallogr. 1997, 53, 240–255.

41. Emsley, P.; Lohkamp, B.; Scott, W. G.; Cowtan, K. Features and development of Coot. Acta Crystallogr., Sect. D: Biol. Crystallogr. 2010, 66, 486–501.

